# High-throughput prediction of protein–protein interactions uncovers hidden molecular networks in biosynthetic gene clusters

**DOI:** 10.1101/2025.10.26.684697

**Authors:** Yoshitaka Moriwaki, Taro Shiraishi, Yohei Katsuyama, Kenichi Matsuda, Toyoyuki Ose, Atsushi Minami, Hideaki Oikawa, Tomohisa Kuzuyama, Ryuichiro Ishitani, Tohru Terada

**Affiliations:** Department of Computational Drug Discovery and Design, Medical Research Laboratory, Institute of Integrated Research, Institute of Science Tokyo, 1-5-45 Yushima, Bunkyo-ku, Tokyo 113-8510, Japan; Department of Biotechnology, Graduate School of Agricultural and Life Sciences, The University of Tokyo, 1-1-1 Yayoi, Bunkyo-ku, Tokyo 113-8657, Japan; Department of Chemistry, Graduate School of Science, Kyoto University, Sakyo-ku, Kyoto, 606-8502, Japan; Collaborative Research Institute for Innovative Microbiology, The University of Tokyo, Bunkyo-ku, Tokyo 113-8657, Japan; Faculty of Pharmaceutical Sciences, Hokkaido University, kita-ku, Sapporo, 060-0812, Japan; Faculty of Advanced Life Science, Hokkaido University, kita-ku, Sapporo, 060-0810, Japan; Department of Chemistry, Institute of Science Tokyo, 2-12-1 O-okayama, Meguro-ku, Tokyo 152-8551, Japan; Division of Chemistry, Faculty of Science, Hokkaido University, Sapporo, 060-0810, Japan

## Abstract

Biosynthetic gene clusters (BGCs) are contiguous genomic regions that encode diverse proteins responsible for natural product biosynthesis. These proteins collectively produce various secondary metabolites with complex chemical structure, including antibiotics and mycotoxins, yet the complete biosynthetic pathways have been experimentally resolved for only a limited number of compounds. Protein–protein interactions within BGCs have recently been recognized as key determinants of intermediate transfer, enzymatic regulation, and structural stability. However, many BGCs still contain proteins of unknown function that cannot be predicted by conventional sequence-based bioinformatics tools, hindering a comprehensive understanding of their biosynthetic pathways. To address this challenge, we built a high-throughput complex prediction pipeline by replacing AlphaFold3’s multiple sequence alignment generation with a faster MMSeqs2. We systematically screened 487,828 protein pairs derived from 2,437 BGCs registered in the Minimum Information about a Biosynthetic Gene cluster (MIBiG) database and predicted 15,438 heteromeric interactions with an ipTM ≥ 0.6. Among them, 1,390 protein pairs exhibited structural homology with an RMSD ≤ 2.0 Å. These predictions highlight interesting molecular mechanisms involving proteins previously annotated as “uncharacterized” or “potentially dysfunctional”. Our analysis further showed that the ipSAE metric can distinguish correct heterocomplex pairs when multiple functionally homologous proteins are present within a BGC. Overall, our computational analysis revealed molecular interaction networks among proteins encoded by each BGC and identified enzyme complexes that are likely functional only when assembled. These predicted complexes may represent previously unrecognized links in their biosynthetic pathways. The complete results are available in a reusable format at https://doi.org/10.5281/zenodo.17451667 to support future experimental validation.

## Introduction

Secondary metabolites are natural products produced by organisms that are not directly involved in their growth, development, or reproduction. Unlike primary metabolites, such as amino acids, nucleotides, and carbohydrates, secondary metabolites often play specialized roles in ecological interactions—such as defense against predators^1^, competition with other organisms^2^, or symbiotic relationships^3^. These compounds are typically produced by biosynthetic gene clusters (BGCs), which are co-located sets of genes that collectively encode enzymes required to produce a specific metabolite and auxiliary proteins such as transporter and regulatory proteins^4, 5^. Secondary metabolites contain a wide variety of chemical structures and classes, including non-ribosomal peptides (NRPs), polyketides (PKs), ribosomally synthesized and post-translationally modified peptides (RiPPs), terpenes, and saccharides, which have attracted significant interest from both enzymology and organic chemistry. Since the 2000s, progress in sequencing technologies, such as next-generation sequencing and metagenomic analysis, along with the development of bioinformatics tools and the expansion of genomic databases, has significantly advanced research on secondary metabolites derived from uncultivable microorganisms. For example, antiSMASH^6, 7, 8, 9, 10, 11, 12, 13^ is a widely used bioinformatics tool for the identification, annotation, and analysis of BGCs in microbial genomes. It also classifies clusters into known types by comparing them to curated reference datasets, including the Minimum Information about a Biosynthetic Gene cluster (MIBiG) database^14^. DeepBGC^15^ incorporates deep learning and natural language processing strategies to capture genomic positional dependencies and order information that are not considered in hidden Markov model (HMM)-based tools such as ClusterFinder^16^ and antiSMASH. This improved the identification of BGCs in bacterial genomes and contributed to the discovery of novel BGCs encoding natural products that could not be detected by experimental methods.

Proteins can expand their functional capabilities by forming complexes with other proteins or ligands. As in other types of gene clusters, the formation of protein complexes plays a crucial role in BGCs. Notably, protein–protein interactions observed in carrier proteins^17^, *trans*-type acyltransferase (AT) polyketide synthase (PKS)^18^, *cis*-type AT PKS^19^, type I/II PKS, and fatty acid synthase^20, 21^ have been extensively studied and discussed. These interactions are key components of the biosynthetic machinery. In homomultimer complexes, residues that are distant in the primary and tertiary structures assemble at the multimer interface to form a single binding pocket^22^. Additionally, some heteromeric complexes influence the stability, specificity, and catalytic activity^23, 24^. Because these interactions underlie the remarkable structural and chemical diversity of the products, many enzymes in these systems have been structurally characterized to clarify their molecular basis to date. Nevertheless, most BGCs still harbor numerous proteins of unknown function, often annotated simply as “hypothetical protein” in MIBiG. Sequence-based bioinformatics tools generally cannot predict the function of these proteins unless homologous proteins with known functions have already been characterized. This limitation represents a major challenge for the comprehensive understanding of their biosynthetic pathways.

Highly accurate prediction of protein tertiary structures, as exemplified by AlphaFold2 (AF2)^25^, has significantly advanced structural bioinformatics. Structures predicted with high confidence are often sufficiently accurate to be used as search models of molecular replacement method of for X-ray crystallography^26, 27^, as well as initial structures for molecular dynamics (MD) simulations and quantum mechanics/molecular mechanics (QM/MM) calculations^28^. Furthermore, AlphaFold provides valuable predictions about protein–protein^29^ interactions that are difficult to obtain from sequence-based bioinformatics tools alone. Notably, AlphaFold predicts inter-residue distances based on co-evolutionary signals concealed in the multiple sequence alignments (MSA) for the query sequence(s). When more than roughly 30–100 effective homologous sequences are available in sequence database, this strategy enables accurate prediction without relying on experimentally determined homologous structures or prior knowledge of protein–protein interactions. One application of protein complex prediction is the discovery and experimental validation of the protein Tmem81, which is conserved across vertebrates during fertilization of sperm and egg but whose function could not be inferred prior to the study^30^.

Here, we report a comprehensive search of all pairwise complex predictions among proteins encoded within each BGC using AlphaFold3 (AF3)^31^. These computational analyses enabled highly sensitive detection of potential homo- and heteromultimer formation that were previously difficult to predict using conventional sequence-based bioinformatics tools. This approach led to the identification of novel pairs of proteins that are likely to act cooperatively. Furthermore, our results suggest that numerous structurally homologous heterocomplexes exist in BGCs and that many of them involve proteins previously annotated as having unknown functions. Finally, we visualized the protein–protein interaction network maps to provide a comprehensive overview in the BGCs. Taken together, these findings offer novel and significant insights into previously uncharacterized proteins in BGCs, thereby providing valuable contributions to both experimental and computational researcher communities engaged in the investigation of secondary metabolites.

## Results

### High-throughput complex prediction pipeline

We developed a computational pipeline for the systematic prediction of protein–protein complexes encoded within BGCs (**Fig. 1**). The pipeline takes as input the GenBank (.gbk) files associated with BGC accession IDs registered in MIBiG. Users may also provide a GenBank files generated by antiSMASH or FASTA-formatted protein sequence files for unregistered BGCs. Here, we focused on 2437 BGCs that are labeled as “active” in MIBiG and targeted proteins with up to 1,950 amino acid residues. For each BGC, complexes were predicted for all possible protein combinations, including self-pairs. Specifically, for a BGC containing *N* qualifying proteins, we performed *N*(*N* − 1)/2 predictions for heteromeric pairs and *N* additional predictions for homomeric pairs, resulting in a total of *N*(*N* + 1)/2 predictions per BGC.

**Fig. 1:**
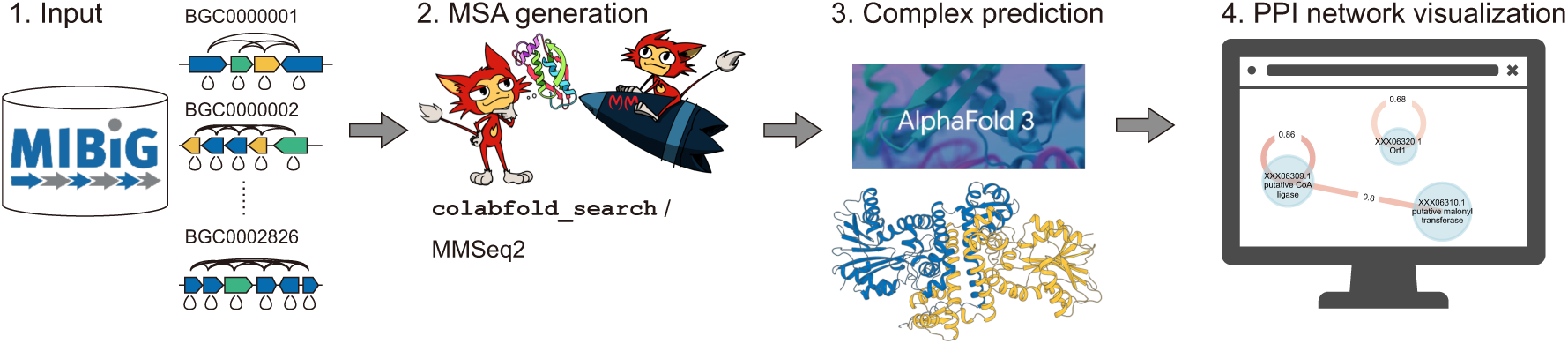
computational workflow for protein complex prediction. 1. Input amino acid sequences. List all pairs of self or other protein sequences encoded in each BGC accession ID, excluding those with a length more than 1950 residues. 2. Generate paired and unpaired MSAs for all the protein pairs using MMseqs2 and *colabfold_search*. 3. Predict the dimeric complex using AlphaFold3. The input JSON files were created with *alphafold3_tools*. 4. Visualize a protein-protein interaction (PPI) map for each BGC accession ID. Output JSON files containing ipTM and ipSAE metrics were also generated.

Highly accurate structure prediction using AF3 requires the generation of multiple sequence alignments (MSAs) for the input sequences. However, comprehensive prediction using the AF3 Web server is not feasible due to its usage limits (currently 30 runs per day). Moreover, the official AF3 inference code available on GitHub is also impractical for the high-throughput workflows, mainly because the MSA generation process using HMMER 3^32^ is slow and cannot be efficiently parallelized. To address these issues, we used *colabfold_search* from LocalColabFold^33^ in conjunction with MMseqs2^34^ to rapidly generate MSAs. The resultant MSAs included both unpaired MSAs for individual proteins and paired MSAs generated by concatenating sequences from the same organism. These were assigned to the “unpairedMsa” and “pairedMsa” keys, respectively, in the AF3 input JSON file. This conversion was performed using the *msatojson* command, which is part of our *alphafold3_tools* toolkit. Using this pipeline, we generated MSAs in 1–2 minutes per complex on a server with 768 GB of RAM or more and efficiently distributed the structure inference tasks for a total of 487,828 protein pairs across multiple GPU-equipped computers.

### Validation of the predicted complexes

We subsequently compared the predicted complexes with the experimentally determined protein complexes deposited in Protein Data Bank (PDB) to evaluate the validity. As of November 30, 2024, a total of 555 homomeric and 30 heteromeric protein entries were found in PDB to exhibit over 95% sequence identity with proteins registered in MIBiG ver. 4.0. Among the 555 homomeric proteins, 408 were annotated as homodimers, and 135 as homotrimers or higher-order assemblies that could be modeled on our GPUs. The remaining 12 entries could not be predicted due to their extremely large complex sizes. A high sequence similarity threshold of 95% was applied to minimize the effects of amino acid differences, insertions, or deletions that could alter the composition of residues at the multimerization interface and thereby lead to inaccurate measurements.

For the protein structures registered in PDB as homodimers, our predicted dimeric models showed ipTM ≥ 0.8 for 294/408 (72.1%) and ipTM ≥ 0.6 for 358/408 (87.7%) (**Fig. 2a**). The ipSAE metric, which represents a corrected ipTM calculated using only interchain residue pairs with reliable PAE scores, correlated well with ipTM above the 0.6 threshold, but approached zero below this cutoff. We next examined the 135 proteins annotated as forming higher-order homomeric complexes. When predicted as dimers, these proteins frequently yielded ipTM values around 0.6. However, when re-predicted with their stoichiometries of biological assemblies registered in PDB, the ipTM values substantially improved, exceeding 0.8 in most cases (**Fig. 2b**). A representative example is CutA1 from *Oryza sativa* (PDB ID: 2ZOM). In the dimeric prediction, the substructure was nearly identical to the crystal structure but exhibited low confidence scores (ipTM = 0.38; ipSAE = 0.001). In contrast, the trimeric prediction, corresponding to the biological assembly, closely reproduced the experimental structure, with improved ipTM and ipSAE values of 0.64 and 0.84, respectively (**Fig. 2c**). These results highlight the limitations of using ipTM or ipSAE thresholds for dimer prediction, especially when the inter-subunit interface is small. Additionally, the ipTM and ipSAE metrics were further improved to 0.93 and 0.86, respectively, when its N-terminal long transit peptide region (residues 1–64) was not included in the prediction. This finding also indicated that the long-disordered region lowered ipTM (**Supplementary Fig. 1**). Despite this, ipSAE remained high, underscoring its reliability as an evaluation metric in such cases.

**Fig. 2:**
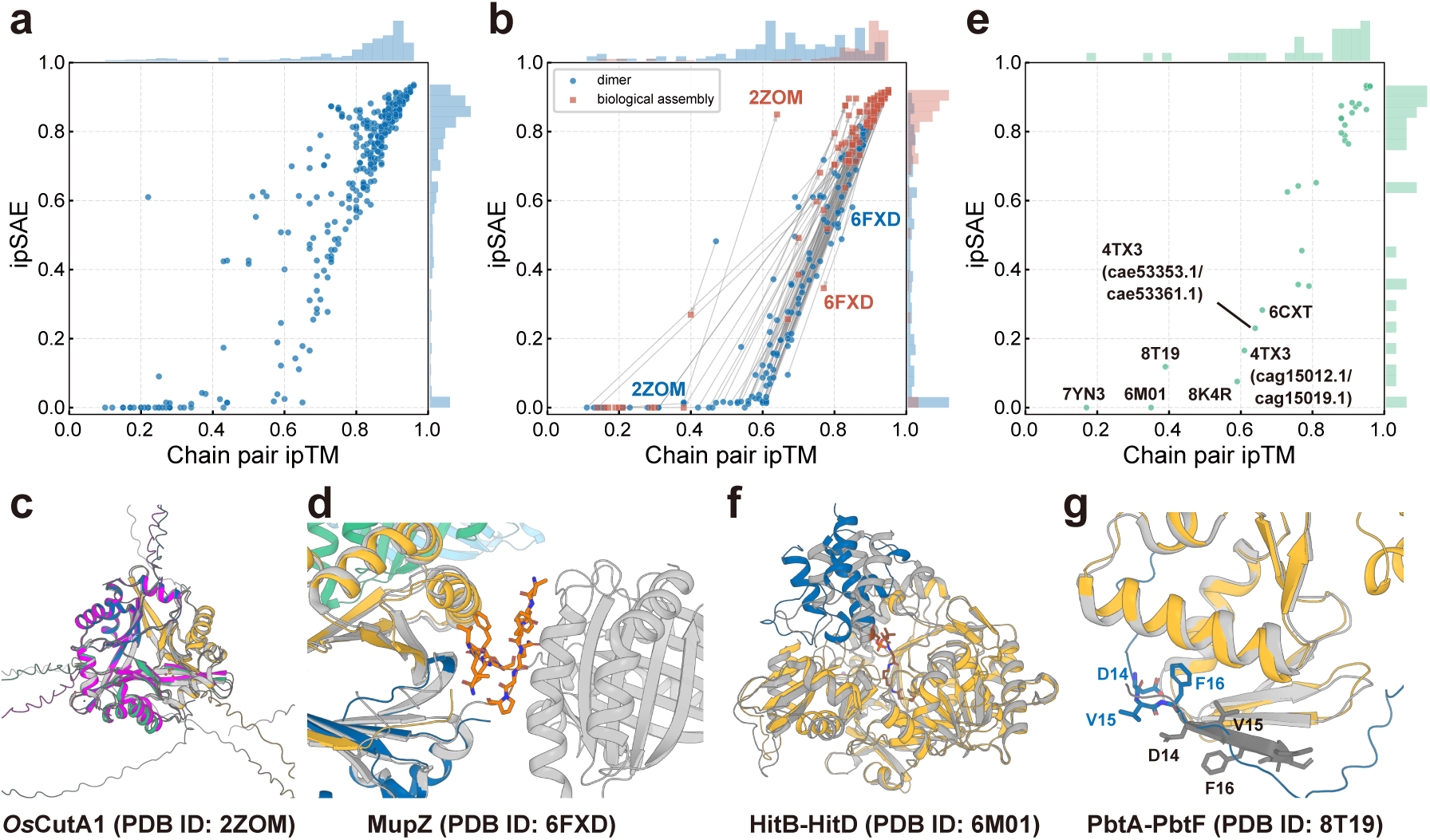
Validation of the predicted complexes. **a** Scatter plot of chain-level ipTM and ipSAE values for predicted homodimers (*n* = 408). Marginal histograms for each axis are displayed above and to the right. **b** Differences in ipTM and ipSAE between the predicted homodimer (blue) and the biological assembly (red). Only biological assemblies forming homomeric complexes larger than dimers and computable on our GPUs are included (*n* = 135). **c** Superimposition of the predicted *Os*CutA1 trimer (GenBank ID: BAT10611.1; blue, yellow, and green) and dimer (purple) with the crystal structure (PDB ID: 2ZOM). **d** Predicted MupZ homotetramer (colored) and its crystal structure (white). The artificial N-terminal peptide (orange) is shown in stick representation. **e** Scatter plot of ipTM and ipSAE values for predicted heterodimers. **f** Predicted HitB (GenBank ID: BAR73008.1)–HitD (BAR73010.1) and its crystal structure (white and gray). The cross-linking probe in the crystal structure is shown in brown stick representation. **g** Superimposition of the predicted PbtA (AGY49587.1; blue)–PbtF (AGY49592.1; yellow) complex with the crystal structure (PDB ID: 8T19; white and gray). Residues of PbtA at the interface are shown in stick representation.

Nevertheless, some of the predicted complexes did not exhibit high ipTM/ipSAE values even when using the stoichiometries of biological assemblies. Closer inspection revealed that several of these cases represented artifacts, either due to misassignments of stoichiometry inferred from the crystal packing using PDBePISA^35^, or the inclusion of artificial sequence tags that induced unnatural oligomerization. The former example is MxaA in *Stigmatella aurantiaca* (GenBank ID: AAK57184.1; PDB ID: 4U7W), which is experimentally validated as a monomer^36^. The latter example is MupZ from *Pseudomonas fluorescens* (**Fig. 2d**; GenBank ID: AFD33556.1; PDB ID: 6FXD), which is registered as a homo-tetramer due to C-terminal His₆-tag–induced tetramerization, although the native protein forms a dimer.^37^ We summarized the proteins exhibiting low ipSAE and ipTM values in **Supplementary Table 1**. These observations highlight the challenges of constructing fully reliable datasets of multimeric assemblies from PDB. Nonetheless, our findings suggest that an ipTM threshold of 0.6 is effective for screening proteins for the potential of homooligomerization.

### Validation of the predicted heteromeric complexes

For the 30 heteromeric complexes, we observed similar ipTM and ipSAE correlations to those in the homomeric complexes. Among these, 6M01^38^, 7YN3^39^, 8K4R^40^, 8T19 (DOI: 10.2210/pdb8T19/pdb), and 4TX3^41^ showed ipTM < 0.6 (**Fig. 2e**). We further investigated the cause of these low ipTM values. First, in PDB entries of 7YN3, 8K4R (VinM-VinL), and 6M01 (**Fig. 2f**), a chemical reagent to induce covalent cross-linking between the two proteins was used to obtain the crystal structure of the heterodimeric complexes. Biologically, one of the proteins in these two PDB entries is an acyl carrier protein (ACP), which is known to form a transient complex with the other protein. Thus, it is speculated that the ipTM does not yield high values for such complexes.

PDB ID 8T19 represents the complex structure of PbtA, a RiPP precursor peptide, and its modifying enzyme PbtF from the GE2270 BGC (BGC0001155). The ipTM and ipSAE values were 0.39 and 0.1, respectively. Interestingly, AF3 predicted that the peptide binds to the β-strand region of PbtF; however, the predicted binding position was shifted by two residues compared with the crystal structure (**Fig. 2g**). We further investigated NisB from *Lactococcus lactis subsp. lactis* and StrB from *Streptococcus thermophilus* LMD-9 (BGC0001209)^42^ to gain insights into proteins that bind RiPP precursor peptides. The crystal structure of NisB in complex with its precursor peptide NisA was not included in the validation dataset because these proteins were fused within the same chain (PDB ID: 4WD9)^43^ or the peptide is chemically modified (PDB ID: 6M7Y)^44^. StrB shares 94.98% sequence identity with its homologous protein SuiB from *Streptococcus suis* (PDB ID: 5V1T)^45^. The predicted complex structures for both matched well with their corresponding crystal structures, showing no residue shift and high ipTM and ipSAE values (**Supplementary Fig. 2**). However, because of the extremely limited number of available structures or experimental data on formation of RiPP precursor peptide–enzyme complexes, a statistical evaluation for them remains difficult.

Taken together, these results indicate that AF3 successfully predicted complex formation (ipTM ≥ 0.6) in 26 out of the 30 heteromeric cases. However, there are limitations in its predictive capabilities, particularly for transient complexes or those with small binding interfaces, such as those formed by ACPs or precursor peptides.

### Novel heterocomplexes

Among 487,828 predicted protein pairs, 15,438 heterodimers exhibited ipTM ≥ 0.6, and 3,754 showed ipTM ≥ 0.8. These include complexes formed by previously uncharacterized proteins. Here, we can hypothesize the functions of these proteins based on the predicted models. For example, a heterodimer of cae45685.1 and cae45686.1 (**Fig. 3a–c**) found in borrelidin BGC from *Streptomyces parvulus* (MIBiG accession ID: BGC0000031)^46, 47, 48^ is annotated as “hypothetical protein” in MIBiG, and to our knowledge, no studies have been reported for these proteins. BLASTp^49^ search predicted both proteins have partial sequence similarity with GCN5-related *N*-acetyltransferase (GNAT). Structure-based searches against PDB100 database using the Foldseek webserver^50^ indicated that the predicted complex structure resembles the single subunit Naa30 of the heterotrimeric NatC complex (PDB ID: 6YGD, chain A)^51^, which catalyzes N-terminal acetylation (Nt-acetylation) of numerous eukaryotic proteins. Superimposition of the complex model on the Naa30 structure revealed that a binding pocket for the cofactor acetyl-CoA is formed between the cae45685.1–cae45686.1 interface, with the acetyl group of acetyl-CoA being exposed to the peptide-binding site of Naa30 (**Fig. 3d**). This suggests that this heterocomplex may also catalyze Nt-acetylation of its substrate. Recently, *Streptomyces mutabilis sp*. MII, which is thought to possess a BGC similar to BGC0000031, has been reported to produce *N*-acetylborrelidin B in addition to borrelidin (**Supplementary Fig. 3a**)^52^. Given that borrelidin B, a known precursor of borrelidin, has an aminomethyl group at C12 carbon, it is plausible that the heterocomplex binds borrelidin B at a peptide-binding site analogous to that of Naa30 and catalyzes its acetylation to yield *N*-acetylborrelidin B (**Supplementary Fig. 3b and 3c**).

**Fig. 3:**
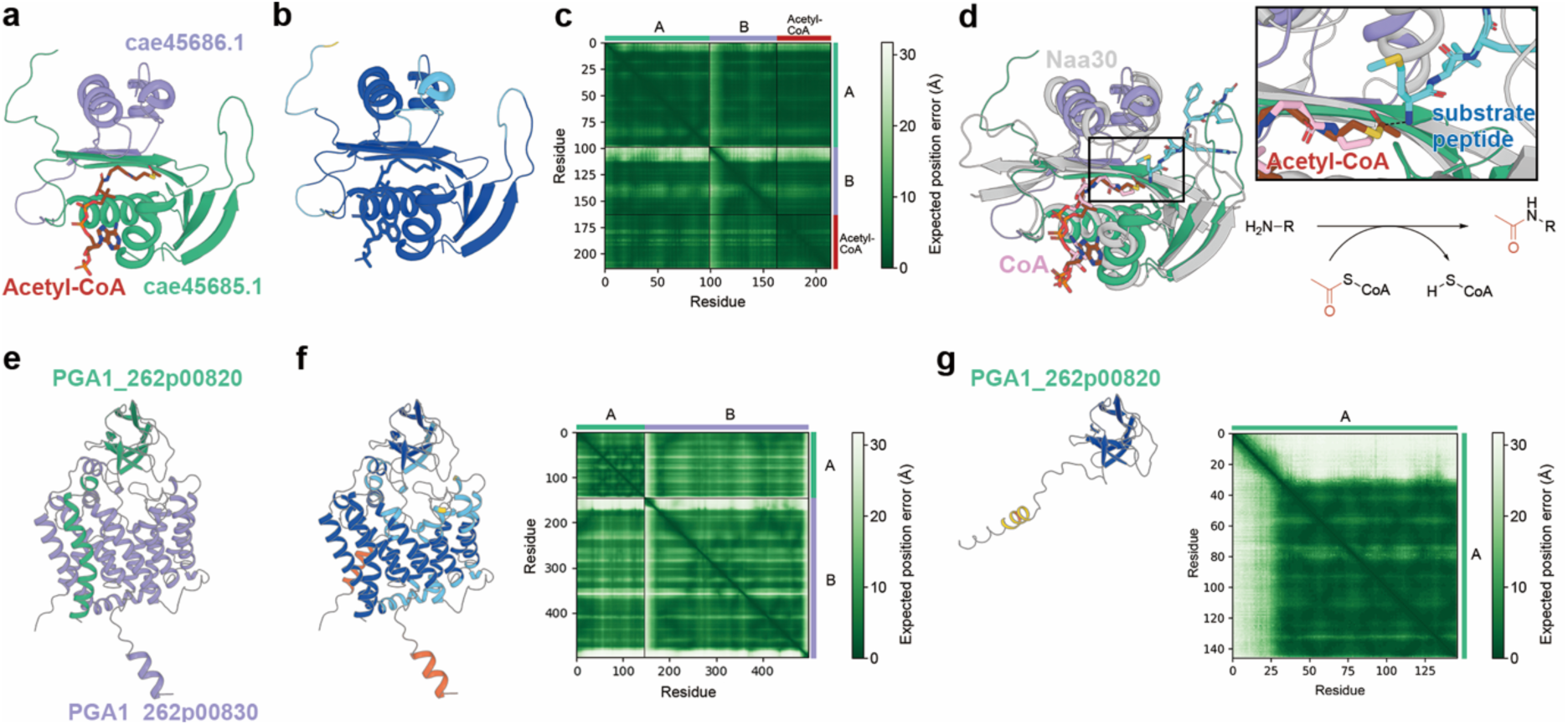
Examples of predicted complex models. **a** cae45685.1 (green)–cae45686.1 (purple)–Acetyl-CoA (brown) trimeric complex model. **b** The predicted complex colored by pLDDT (orange, 0–50; yellow, 50–70; cyan, 70–90; and blue, 90–100). **c** The predicted aligned error (PAE) matrix with chain coloring of Panel **a** on the side bars. The dashed black lines indicate the chain boundaries. **d** Superimposition of the Naa30 crystal structure (white) on the predicted model shown in Panel **a**. The N-terminal region of major capsid protein as a substrate peptide (Uniprot ID: P32503) is shown as cyan stick. **e** PGA1_262p00820 (green)–PGA1_262p00830 (purple) complex model found in BGC0000932. **f** pLDDT-colored predicted PGA1_262p00820–PGA1_262p00830 complex. The PAE matrix with chain coloring of Panel **f** is shown on right panel. **g** Predicted PGA1_262p00820 monomer structure colored by pLDDT. The right panel shows the PAE matrix. These data were obtained from the AlphaFold Protein Structure Database^53^ (UniProt ID: I7E6A3).

We also identified several Domain Unknown Function (DUF) proteins whose function appears to arise only upon complex formation. For example, PGA1_262p00820 (Genbank ID: AFO93364.1) in tropodithietic acid BGC from *Phaeobacter inhibens* DSM 17395 (MIBiG accession ID: BGC0000932)^54^ was annotated as a DUF4399 domain–containing protein, a family for which functional characterization remains scarce. Our analysis predicted that this protein forms a complex with PGA1_262p00830 (AFO93365.1), a DoxX-family protein, with an ipTM of 0.82 and an ipSAE of 0.64 (**Fig. 3e**). Notably, the N-terminal region of PGA1_262p00820 was predicted to be disordered as a monomer; however, in the heterodimer, this region adopted an α-helical conformation with high pLDDT and formed an interaction interface with a helix of PGA1_262p00830 (**Fig. 3f and g**). Given that DoxX-family proteins are localized to the cell membrane, PGA1_262p00820 is expected to be embedded with PGA1_262p00830 in the cell membrane. Thus, the two proteins may act cooperatively as membrane-bound enzymes or transporters.

### Structurally homologous heterocomplexes

By examining the predicted heterodimers, we identified 952 pairs (4.9%) in which the structural RMSD between constituent proteins was ≤1.0 Å, and 1390 pairs (7.1%) with RMSD ≤ 2.0 Å (**Fig. 4a**). The median ipTM and ipSAE values for the protein pairs with RMSD ≤2.0 Å were 0.82 and 0.71, respectively, indicating that these predictions were made with high confidence. This observation suggests that structurally homologous heterodimers are present across BGCs and contribute to the biosynthesis. A well-characterized and popular example is a ketosynthase (KS)– chain length factor (CLF) complex, which constitutes the central catalytic machinery in Type II PKSs. In this complex, the KS subunit catalyzes the decarboxylation of the malonyl unit tethered to a phosphopantetheinyl arm of an ACP, and the condensation of the resulting enolate anion with a growing polyketide chain attached to its catalytic cysteine. The CLF subunit is a KS homolog lacking catalytic residues, but the internal tunnel formed by the complex plays a critical role in determining the polyketide chain length. The actinorhodin polyketide beta-ketoacyl synthase alpha (GenBank ID: CAC44200.1) and beta (CAC44201.1) subunits of BGC0000194 have been identified as KS and CLF, respectively, and their heterocomplex structure was determined (PDB ID: 1TQY)^55^. The protein pair was predicted to form a complex with high ipTM and ipSAE values of 0.96 and 0.93, respectively, and closely matched the crystal structure (**Fig. 4b**). Interestingly, despite the KS and CLF subunits sharing the same fold with an RMSD of 1.02 Å, the homodimeric prediction for KS yielded relatively low scores (ipTM = 0.65, ipSAE = 0.38), and CLF showed almost no signal (ipTM = 0.20, ipSAE = 0.0). These findings indicate that AF3 is capable of correctly identifying heterocomplex formation independent of overall structural similarity between monomers.

**Fig. 4:**
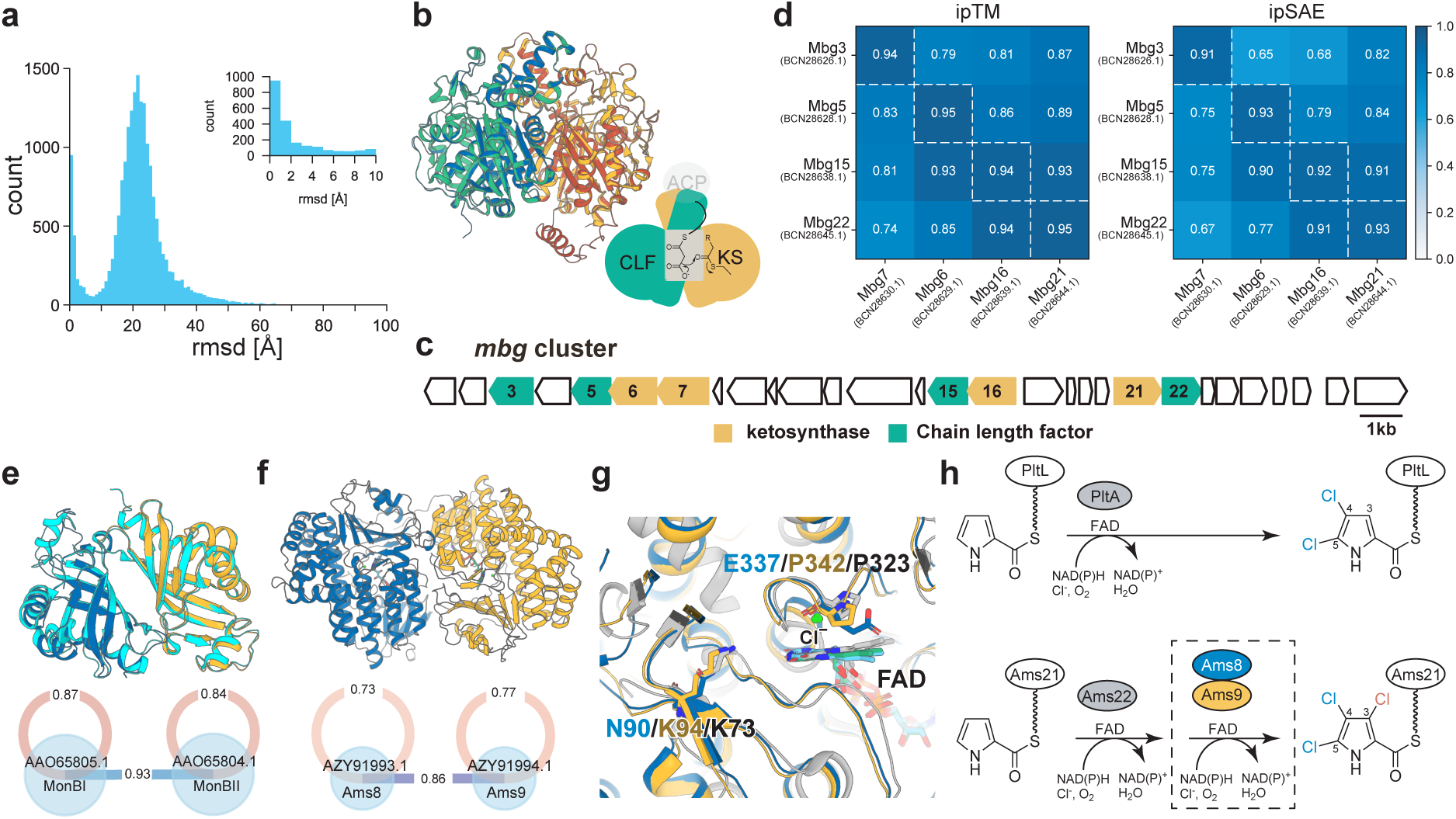
Predicted structurally homologous complexes. **a** Histogram of structural RMSD values between monomeric units of predicted heterodimer pairs. An inset shows a magnified view of the 0–10 Å range. **b** Superimposition of the predicted complex structure of actinorhodin polyketide β-ketoacyl synthase alpha (KS) and beta (CLF) subunits (blue and red) on the crystal structure (PDB ID: 1TQY; green and yellow). KS is shown in blue and green, and CLF in red and yellow. **c** *mbg* cluster. Only the genes encoding KS and CLF are depicted. **d** Computed ipTM and ipSAE values for each complex prediction of the four KS (horizontal) and four CLF (vertical) subunits in *mbg* cluster. **e** Superimposition of the predicted MonBI (blue)–MonBII (yellow) heterodimer on the crystal structure of the MonBI homodimer (PDB ID: 3WMD; cyan). The computed ipTM values for each homo- and hetero-dimer are shown below. **f** Predicted Ams8 (blue)–Ams9 (yellow) complex. **g** Close-up view of the FAD binding site in the predicted structures of Ams8 (blue) and Ams9 (yellow). The pltA crystal structure (PDB ID: 5DBJ) is shown in gray. Residues presumed to be important for chlorination activity are shown in stick representation. **h** Proposed trichlorination mechanism in the armeniaspirol BGC (bottom). Formation of the dichloropyrrole moiety catalyzed by PltA during pyoluteorin biosynthesis is shown on the top for comparison.

Subsequently, we examined whether ipTM and ipSAE could identify correct KS–CLF pairs within a BGC cluster encoding multiple KS and CLF proteins. In our previous work, we experimentally validated the correct pairing in *mbg* cluster (BGC0002441; **Fig. 4c**) as Mbg7–Mbg3, Mbg6–Mbg5, Mbg16–Mbg15, and Mbg21–Mbg22 ^56^. Our all-by-all complex predictions for the 4 × 4 combinations showed that both ipTM and ipSAE values reached their maximum values for the experimentally validated pairs (**Fig. 4d**). Notably, ipSAE yielded lower values for incorrect pairings than ipTM, indicating greater discriminatory power. A similar trend was observed in the Ysf cluster (BGC0002547)^57^, which encodes three KSs and three CLFs, where the highest values again corresponded to the experimentally confirmed pairings (**Supplementary Fig. 4**). Collectively, these results demonstrate that the ipTM and ipSAE metrics provide reliable indicators for identifying true KS–CLF pairs and predicting their released polyketide intermediates, even when the corresponding genes are not located adjacent within the BGC.

Our comprehensive predictions further identified 381 protein pairs that are likely to form both homodimers and heterodimers, meeting the criteria of ipTM ≥ 0.6 and ipSAE ≥ 0.6. A representative example is the pair MonBI and MonBII from the monensin BGC (BGC0000100). Although the crystal structure of the MonBI homodimer has been determined in our previous work (PDB ID: 3WMD)^58^, biological assays have shown that MonBI also forms a heterodimer with MonBII, and this complex displays enzymatic activity. AF3 predictions indicated that MonBI and MonBII form homodimers with ipTM values of 0.87 and 0.84, respectively, while they form a heterodimer with a higher ipTM value of 0.93 (**Fig. 4e**). The predicted MonBI–MonBII heterodimer structure closely matched the MonBI homodimer crystal structure, with an RMSD of 1.0 Å. This result suggests that dimer formation between these two proteins is competitive, and that the heterodimer is preferentially formed.

Proteins predicted to form both homo- and hetero-dimers were also observed among other functional proteins. We highlight three flavin-dependent halogenase-like proteins—Ams8, Ams9, and Ams22—from the armeniaspirol BGC (BGC0002022)^59^ as an example. Within the BGC, Ams21 and Ams22 share sequence similarities of 43% and 65% with PltL and PltA, respectively, from the characterized pyoluteorin BGC (BGC0000127), suggesting that Ams21 functions as an ACP and Ams22 as a flavin-dependent halogenase. Moreover, the structure of the active site of Ams22 is highly similar to that of PltA (PDB ID: 5DBJ)^60^, which catalyzes the 4,5-dichlorination of the pyrrolyl moiety covalently attached to the phosphopantetheinyl arm of PltL, strongly suggesting that Ams22 catalyzes the same reaction (**Supplementary Fig. 5**). However, unlike in the pyoluteorin BGC, the armeniaspirol BGC was reported to produce a 3,4,5-trichlorinated pyrollyl intermediate during the biosynthetic process^59^. Thus, the remaining halogenase-like enzymes, Ams8 and Ams9, are hypothesized to catalyze chlorination at the 3-position of the pyrrolyl moiety, although the mechanism remains unclear^59^.

Our structure prediction pipeline showed that Ams8 and Ams9 may form homodimers with ipTM values of 0.77 and 0.73, respectively, whereas their heterodimer displayed a higher ipTM value of 0.86 (**Fig. 4f**). Intriguingly, the predicted structure of Ams8 revealed that E337 is positioned at a site that would inhibit the binding of flavin adenine dinucleotide (FAD), an essential cofactor for halogenation. In addition, the conserved lysine residue required to mediate the reaction between the substrate and hypochlorous acid^61, 62, 63^ generated from FAD, molecular oxygen, and chloride ions—is replaced by N90 in Ams8. (**Fig. 4g**). These unusual residue substitutions strongly suggest that Ams8 lacks halogenase activity. By contrast, Ams9 and Ams22 retain these conserved residues. However, gene deletion of *ams8* completely abolished the biosynthetic process of armeniaspirol, whereas deletion of either *ams9* or *ams22* reduced its production by 100-fold but did not lead to complete loss^59^. This indicates that Ams8 is more essential for the biosynthetic pathway. Overall, these findings suggest that Ams9 functions as the enzyme catalyzing trichlorination of 4,5-dichloropyrrolyl–*S*-Ams22, while Ams8 acts as an auxiliary protein that supports the activity of Ams9 through complex formation (**Fig. 4h**).

### Visualization of the protein–protein interaction network map

To facilitate the use of our computational results by a broad range of researchers, we created a web site that visualizes protein–protein network diagrams for the 2437 BGCs (http://vivace.bi.a.u-tokyo.ac.jp/network_in_bgc/publish.html). On this site, users can filter the data by MIBiG accession ID, taxonomy, class, or compound names, and view both the network diagrams and representative chemical structures of the products (**Fig. 5a**). Proteins are represented as nodes, and pairs of proteins predicted to form homodimers or heterodimers are connected by edges when they exceed a certain threshold (**Fig. 5b**). The JSON files and Python scripts used to generate these network diagrams are available in the supplementary files, allowing users to perform their own analyses.

**Fig. 5:**
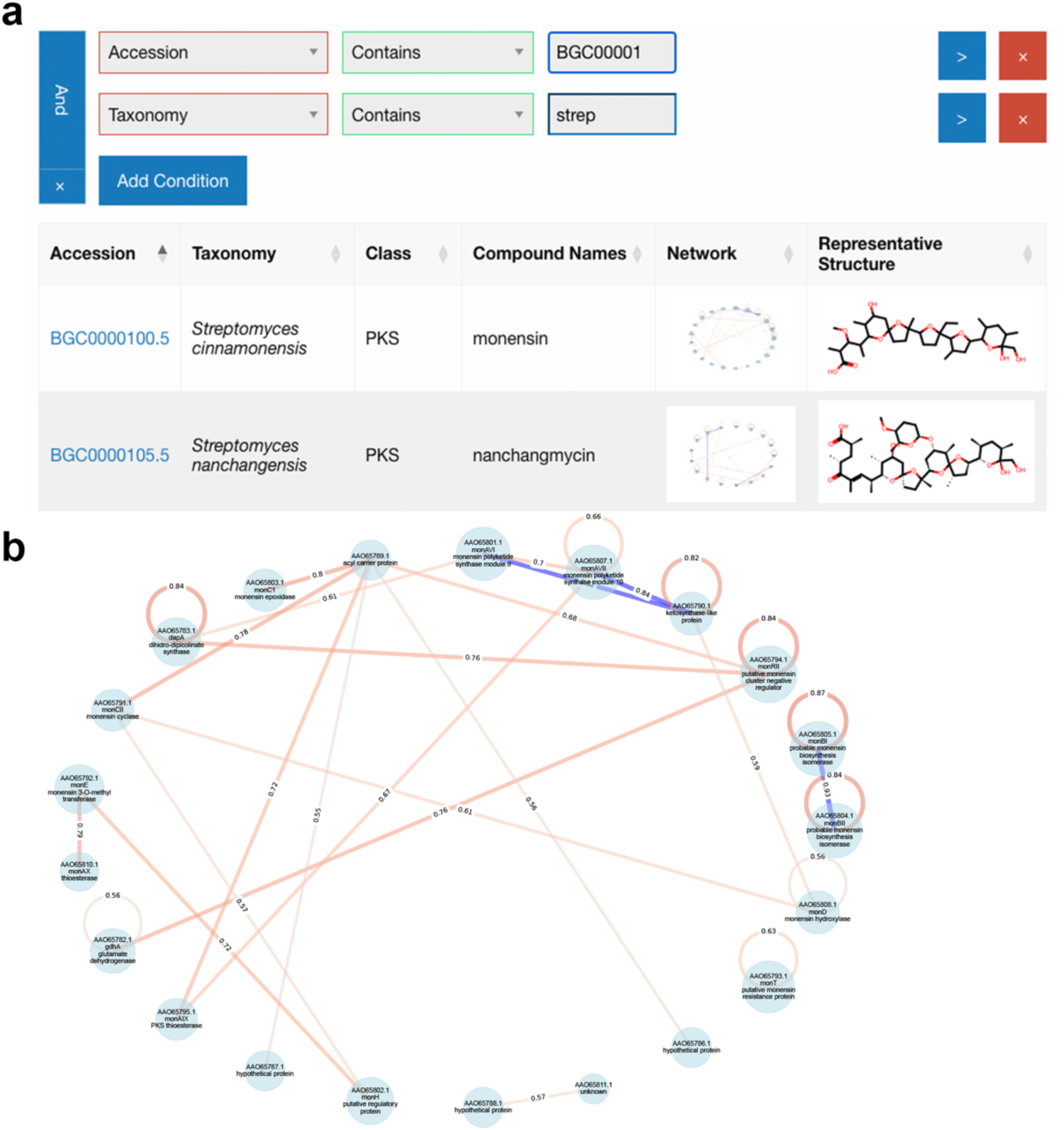
Visualization of the protein–protein interaction (PPI) network map. **a** The interface on the website. **b** The PPI network map in the monensin BGC (BGC0000100). The nodes represent proteins encoded within the BGC. Edges indicate predicted interactions, with labels showing the corresponding ipTM values. Blue edges denote structurally homologous interactions. Only nodes connected by edges above the ipTM threshold are displayed (ipTM > 0.55 was applied on the website).

## Discussion

One of the core challenges in the field of secondary metabolism and biosynthesis research is to elucidate the origins and biosynthetic mechanisms of the vast repertoire of natural products. In this study, we established a high-throughput complex prediction pipeline based on AF3 and demonstrated that it enables comprehensive detection of protein–protein interactions within BGCs. We further showed that BGCs contain a considerable proportion of protein homo- or hetero-complexes whose function emerges only upon complex formation, as well as structurally homologous heterocomplexes.

Although our pipeline and network maps provide valuable insights into the interactome in BGCs, they have three limitations. First, AF2 (including AlphaFold-Multimer) and AF3 often fail to accurately predict weak or transient protein complexes. This limitation arises because their algorithms infer three-dimensional protein structures primarily from coevolutionary signals in the MSA derived from individual amino acid sequence(s), and inter-chain interactions were not considered during their training processes^31^. Carrier proteins, such as ACPs in PKSs and peptidyl carrier proteins (PCPs) in NRPSs, exemplify this limitation. These proteins tether biosynthetic intermediates via a thioester bond to the thiol group of a phosphopantetheinyl arm attached to a conserved serine residue. They then interact with multiple catalytic domains in a specific yet transient manner to promote stepwise transformations of the substrate. Although identifying their partner enzymes is crucial for studying biosynthetic pathways, improving their prediction accuracy remains an important technical challenge. Similarly, predicting the binding position of precursor peptides in RiPPs is also difficult for the same reason.

Second, our computational pipeline did not perform complex predictions for proteins larger than 1,950 residues due to computational limitations. Such large proteins are typically found in type I PKS or NRPS systems, where multiple catalytic domains are linked within a single polypeptide to form the core scaffold of the product. While these modular proteins mainly assemble the product through their intramolecular domain–domain interactions, some tailoring enzymes within the same BGC can modify intermediates tethered to the modular proteins, thereby expanding the chemical diversity of the final products.^64, 65^ Prior prediction of these interactions could help identify novel tailoring enzymes responsible for introducing unusual functional groups; however, such predictions were not fully performed in this study.

Third, our method may fail to correctly predict the complex formation of proteins with potentially conformational changes. Although talaromyolides *tlx* BGC from *Talaromyces purpureogenus* has not been registered in the current MIBiG dataset yet, the TlxJ–TlxI heterodimer crystal structure (PDB ID: 7VBQ)^66^ is notable case to discuss the issue (**Supplementary Fig. 6a**). While sequence analysis suggests that both TlxJ (GenBank ID: BDC03472.1) and TlxI (BDC03471.1) were nonheme iron oxygenases and the RMSD between the two proteins was 1.6 Å, TlxI is suggested to be enzymatically dysfunctional as oxidases due to the lack of an arginine residue for α-ketoglutarate (α-KG) binding and serve as an auxiliary protein to assist in TlxJ expression enhancing its catalytic activity^66^. The gel filtration analysis and the X-ray crystallography indicated that TlxI is monomeric (PDB ID: 7VBR).^66^ However, while TlxJ–TlxI was correctly predicted as a heterocomplex with an ipTM of 0.91 (**Supplementary Fig. 6b**), TlxI was predicted to form a homodimer with an ipTM of 0.85 (**Supplementary Fig. 6c**). Interestingly, the crystal structures revealed that TlxI undergoes a large conformational change when forming a heterocomplex with TlxJ. Yet the predicted TlxI structure was close to the conformation observed in the heterocomplex crystal structure in both monomeric and homomeric forms (**Supplementary Fig. 6c and d**). This observation is consistent with the caveat that AF3 does not adequately predict proteins that undergo conformational changes^31^. By contrast, AF3 successfully predicted the homodimer of MonBI with an ipTM of 0.87 and the heterodimer with MonBII with a higher ipTM of 0.93 (**Fig. 4e**). These results suggest that MonBI does not undergo significant conformational changes upon dimerization. To our knowledge, MSA subsampling with AF2 or ColabFold^67^ is one of the best strategies for predicting such conformational changes in advance. However, we did not apply this approach in this study because of its high computational cost.

Although several technical challenges remain in accurately predicting protein complexes, our comprehensive approach uncovered numerous plausible complexes hidden within BGCs. Notably, the structurally homologous heterodimers identified in this study may serve as intriguing targets for future research, particularly in the context of microbial evolution. According to the genome streamlining theory, natural selection in prokaryotes tends to eliminate genetic redundancy and nonessential sequences, leading to the evolution of highly compact and efficient genomes^68, 69^. Thus, the structurally homologous complexes are unlikely to be mere products of gene duplication but are instead expected to possess distinct functional roles such as regulation or assistance in protein expression. Examples of the former function have been reported in both homo-^70, 71^ and hetero-complexes^66^, and their characterization is relatively straightforward. In contrast, the latter function can only be uncovered through co-expression of a specific protein combination, and its underlying molecular mechanisms remain largely unexplored. Among the reported cases, MonBI–MonBII^58^ and LstA–LstB^72^ become functional only when co-expressed. Co-expression of LxmK and LxmY has been shown not only to improve stability but also to increase yield.^73^ Interestingly, TalA from *Talaromyces pinophilus* LD-7^74^ and Lsd19 from *Streptomyces lasaliensis*^75, 76, 77, 78^ have been identified as naturally occurring chimeric enzymes, in which two structurally homologous protein domains are fused into a single polypeptide chain. Such protein (or domain) pairs may exist to expand the range of catalytic functions that can be achieved by a single enzyme alone.

Currently, sequence-based prediction tools such as DeepBGC and ClusterFinder are widely used to identify BGCs that produce novel natural products. In contrast, structure-based approaches have the potential to elucidate the molecular mechanisms underlying these products based on the relationship between protein function and tertiary structure. The development of prediction tools for protein structures and their interaction networks is a key step toward unraveling enzymatic mechanisms and entire metabolic pathways, even in uncultivable microorganisms. Our results provide a comprehensive overview of the constituent proteins within BGCs and highlight intriguing protein complexes for future investigation. These bottom-up approaches will accelerate identification of protein pairs that should be co-expressed or co-analyzed and deepen our biochemical understanding, ultimately providing insights essential for the rational engineering of enzymes.

## Methods

### Structure Prediction Workflow

#### Input amino acid sequences

We selected the proteins derived from BGCs for comprehensive protein-protein complex prediction based on the following criteria. First, we focused on 2,437 “active” BGC accession IDs deposited in MIBiG version 4.0 (2024-11-15) that are currently maintained and updated. IDs marked as “retired” or “pending” were excluded from analysis. Second, among the proteins encoded in these BGCs, those with more than 1,950 amino acids, which are commonly found in PKS- or NRPS-modular type proteins, were excluded from the prediction, as they can exceed the computational limits of modeling using AF3. The amino acid sequences were obtained from the GBK format files provided by MIBiG. Using the amino acid sequences that meet the criteria, we performed exhaustive predictions of homo- and heterodimeric complexes for each BGC.

#### MSA generation and structure prediction

Despite recent advances, protein structure prediction with AF3 still relies on sufficient number of multiple sequence alignments (MSAs) for the query sequences. To achieve both high accuracy and high-throughput prediction, we replaced the default MSA generation pipeline using HMMER 3^32^ implemented in AF3 with a faster alternative: the *colabfold_search* command in LocalColabFold^33^. MMseqs2 version 15-6f452^34^ was used to build two sequence databases, Uniref30_2302 and colabfold_envdb_202108 (released on July 31, 2023), which can be downloaded from https://colabfold.mmseqs.com/ and generate MSAs for the query sequences. The obtained MSAs have been shown to be sufficient for producing highly accurate predicted models with AF2 or ColabFold in most cases.^33^ In addition to unpaired MSAs, we also obtained paired MSAs to further improve the accuracy of complex modeling. The protein template search against PDB100 database was not performed, as it is known to improve monomeric but not complex prediction accuracy. The MSA generation was performed on the SuperComputer Flow Type III subsystem at Nagoya University. This system comprises 16 Intel Xeon Platinum 8280M processors (2.7 GHz, 28 cores each) and 24 TiB of RAM per node. Each query took approximately 1–2 minutes on the calculation node. The obtained a3m-formatted MSA files were added to the “pairedMsa” and “unpairedMsa” keys of the AF3 input JSON file using the *msatojson* command of the alphafold3_tools software that we developed and made available on our GitHub repository: https://github.com/cddlab/alphafold3_tools/. The “modelSeeds” value in the AF3 input JSON file was set to 1 and five models were generated for each prediction. For dimer with a total number of residues below 1500, the complex structure prediction was performed using the SuperComputer Flow Type II subsystem with NVIDIA GPU V100; for those with 1500 to 2000 residues, our in-house NVIDIA GPU RTX4090; and for those with 2000–3900 residues, NVIDIA GPU H100 (VRAM 94GB). The Predicted Aligned Error (PAE) of the predicted structures was visualized using the *paeplot* command of alphafold3_tools.

### Validation of the predicted complex structures

#### Metrics

To identify biologically plausible protein–protein interactions in the AF3-predicted complexes, we employed two metrics: the interface predicted TM-score (ipTM) and the interaction prediction score from aligned errors (ipSAE)^79^. The ipTM measures the accuracy of the predicted relative arrangement of subunits. Values ≥ 0.8 indicate confident and high-quality predictions, < 0.6 suggest failure, and 0.6–0.8 represent an uncertain range.^29, 31^ The ipTM values are provided in the AF3 output JSON files. The ipSAE is a modified scoring function that evaluates the quality of predicted interactions by focusing on residue pairs likely to participate in the interface, rather than the entire sequence. This metric can be computed by applying the ipSAE script^79^ to the AF3 output JSON file with both the PAE cutoff (pae_cutoff) and Cα-Cα distance cutoff (dist_cutoff) set to 10. We did not use other metrics such as pDock^80^, pDockQ2^81^, or LIS^82^ in this analysis, although they serve similar purposes in evaluating protein–protein interactions.

#### Validation datasets and complex analysis

We evaluated both ipTM and ipSAE metrics for the predicted homo- and heterodimeric structures by validating them against experimentally determined complexes. For proteins encoded in the BGCs listed in MIBiG version 4.0 and also registered in the Protein Data Bank (PDB), we retrieved their biologically relevant complex structures (also referred as “biological assemblies”) and stoichiometry information were retrieved from Protein Data Bank (PDB) as of November 30, 2024. To minimize the impact of amino acid substitutions on protein oligomerization, the sequence identity threshold was set to 95%. Additionally, for complexes whose biological assemblies contained more than two chains and did not exceed 3,900 amino acid residues in total, we performed further structure predictions with the stoichiometries to obtain the corresponding ipTM and ipSAE metrics.

For the obtained heterodimeric structures, the RMSD of the backbone heavy atoms (N, Cα, C, and O) between the two subunits was calculated using Gemmi version 0.7.3 (https://github.com/project-gemmi/gemmi) ^83^, with the maximum number of outlier rejection cycles set to 5. To prevent calculation errors caused by very short sequences (e.g., precursor peptides), RMSD values were not computed for proteins with fewer than 20 residues.

### Network visualization

The protein–protein networks obtained for the 2437 BGCs were drawn using NetworkX version 3.5. Each protein was represented as a node, and edges were drawn between heterodimers that showed ipTM > 0.55 in the AF3 structural prediction. Proteins that formed homodimers at the same threshold were represented with self-loop edges. Then, the network maps were output in SVG image format and made available on a web server. Additionally, the JSON files and Python codes used for generating these network maps will be published as supplementary files, allowing users to regenerate them in their own environments using arbitrary thresholds.

## Data availability

The predicted protein–protein interaction network and JSON files underlying this article are available in Zenodo at https://doi.org/10.5281/zenodo.17451667. The data used for the plots in **Fig. 2** are provided as a Supplementary Excel file.

## Code availability

The python scripts used for the preprocessing, evaluation, and visualization are publicly available at https://github.com/cddlab/bgccomplexbuilder. The *msatojson*, *paeplot*, and other utility commands to process the input and output files of AlphaFold3 can be downloaded as *alphafold3_tools* (https://github.com/cddlab/alphafold3_tools).

## Acknowledgements

This work was supported by the Ministry of Education, Culture, Sports, Science and Technology, Japan [Grant-in-Aid for Transformative Research Areas, 22H05126 to Y.M. and T.T., 22H05120 to T.K. and T.S., 22H05130 to Y.K., 22H05128 to K.M., 25H01575 to A.M., and 25H02250 to R.I.; Grant-in-Aid for Scientific Research (JSPS KAKENHI), 25K01890 to Y.M. and 19H04634 to T.O.]. Additional support was provided by the Medical Research Center Initiative for High Depth Omics at the Institute of Science Tokyo, Nanken-Kyoten at the Institute of Science Tokyo 2025, and the Multilayered Stress Diseases project (JPMXP1323015483) at the Institute of Science Tokyo. This work utilized computational resources from the supercomputer “Flow” provided by the Information Technology Center at Nagoya University through the HPCI System Research Project (Project IDs: jh240001 and jh250002) and the TSUBAME 4.0 supercomputer at the Institute of Science Tokyo.

We thank Dr Naruki Yoshikawa for a fruitful discussion.

## Author information

### Contributions

Yoshitaka Moriwaki: Conceptualization [lead], Data curation [lead], Formal analysis [lead], Funding acquisition [equal], Investigation [lead], Methodology[lead], Resources [equal], Software [lead], Validation [lead], Visualization [lead], Writing—original draft [lead], Writing—review & editing [equal]), Taro Shiraishi: Conceptualization [equal], Investigation [support-ing], Resources [equal], Validation [equal], Writing—review & editing [equal], Yohei Katsuyama: Conceptualization [equal], Investigation [supporting], Validation [equal], Writing—review & editing [equal], Kenichi Matsuda: Conceptualization [equal], Investigation [supporting], Writing—review & editing [equal], Toyoyuki Ose: Writing—review & editing [supporting], Atsushi Minami: Writing—review & editing [supporting], Hideaki Oikawa: Writing—review & editing [supporting], Tomohisa Kuzuyama: Funding acquisition [lead], Project administration [lead], Writing—review & editing [equal], Ryuichiro Ishitani: Funding acquisition [equal], Resources [equal], Supervision [equal], Writing— review & editing [equal], Tohru Terada: Funding acquisition [equal], Project administration [lead], Resources [equal], Supervision [equal], Writing—review & editing [equal]

## Ethics declarations

### Competing interests

The authors declare no competing interests.

## Notes

https://doi.org/10.5281/zenodo.17451667

### Competing Interest Statement

The authors have declared no competing interest.

### Summary of Updates

Title and Introduction section updated to clarify the significance.

