## Supplementary File 1 for "High-throughput prediction of protein–protein interactions uncovers hidden molecular networks in biosynthetic gene clusters"

**Supplementary Table S1: Proteins exhibiting low ipTM and ipSAE values in our prediction pipeline.**

| MIBiG accession ID | GenBank ID | PDB ID[a] | Stoichio-metry registered in PDB | ipSAE (2-mer) | ipTM (2-mer) | ipSAE (biological assembly) | ipTM (biological assembly) | Notes |
|---|---|---|---|---|---|---|---|---|
| 0000294 | BAC87903.1 | 6H7F, 6H7V, 6HCP | 3-mer, monomer, 3-mer | 0 | 0.16 | 0 | 0.17 | Biophysical methods (gel filtration and light scattering) do not support a trimeric assembly.[1] |
| 0000610 | ACR48330.1 | 4ZA1 | 3-mer | 0.82 | 0.87 | 0 | 0.38 | Global Stoichiometry: Homo 3-mer, Local Stoichiometry: Homo 2-mer. |
| 0000668 | AHI58816.1 | 6GEM | 4-mer | 0 | 0.28 | 0 | 0.3 | Biological Assembly Evidence: gel filtration |
| 0000898 | AAC68683.1 | 8HZV, 8HZY | 4-mer, monomer | 0 | 0.23 | 0 | 0.29 | The C-terminal His6-tag of PDB ID: 8HZV may cause the multimerization. 8HZY is not. |
| 0000923 | AAC21672.1 | 3E59, 3EAT, 4YLM | Monomer, 3-mer 3-mer | 0 | 0.21 | 0 | 0.19 | Analyzed by size-exclusion chromatography and eluted midway between the expected times for a dimer and a trimer (data not shown).[2] |
| 0000929 | ABQ88342.1 | 5JDY, 5JDZ, 5JE0, 5JE1, 5JE2, 5JE3, 5JE4, 5JE5, 5JE6 | 4-mer | 0.012 | 0.38 | 0 | 0.2 | The authors conclude that the complexes are most likely to be monomeric.[3] |
| 0001738 | BBC43190.1 | 7DMN, 7DMO, 7E5T, 7E5U, 7E5V | 6-mer | 0 | 0.46 | 0 | 0.21 | Biological Assembly Evidence: gel filtration.[4] |
| 0002109 | QDJ74280.1 | 6J31, 6J32 | 3-mer | 0 | 0.15 | 0 | 0.16 | No description |
| 0002494 | AAD48879.1 | 1L5A | 3-mer | 0 | 0.13 | 0 | 0.14 | VibH is a monomer consisting of two domains.[5] |
| 0002681 | CAG44663.1 | 5NBC, 5NHK | 4-mer | 0.48 | 0.47 | 0.27 | 0.4 | Biological Assembly Evidence: light scattering, gel filtration, SAXS |

[a] The underlined PDB ID was used to compute the ipSAE and ipTM metrics.

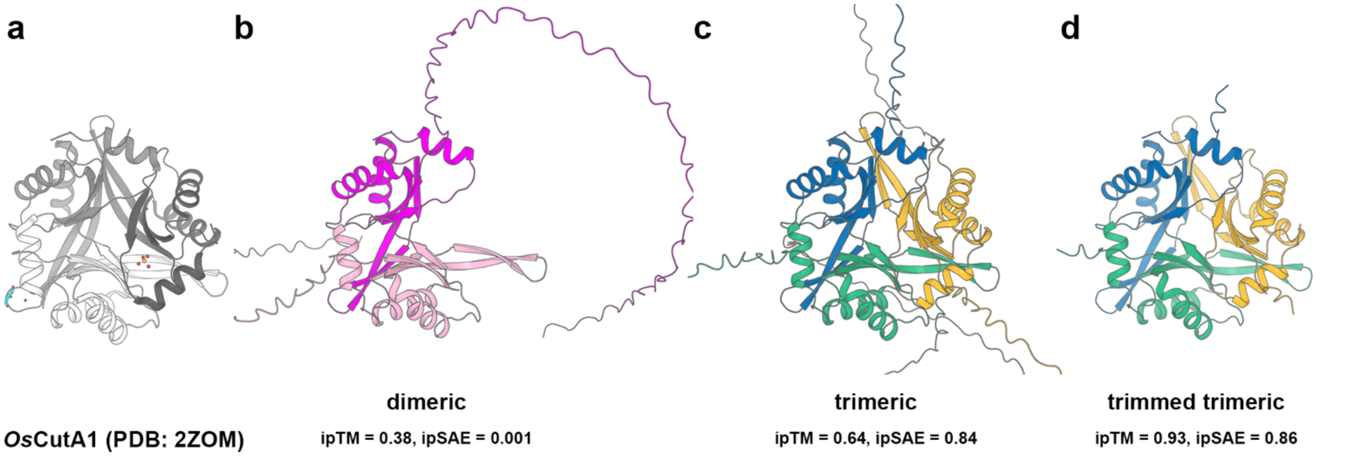


**Supplementary Fig**. **1: Structural comparison of *Os*CutA1. a** Crystal structure (PDB ID: 2ZOM). **b** Predicted dimer structure. **c** Predicted trimer structure. **d** Predicted trimer structure using residues 65–177.

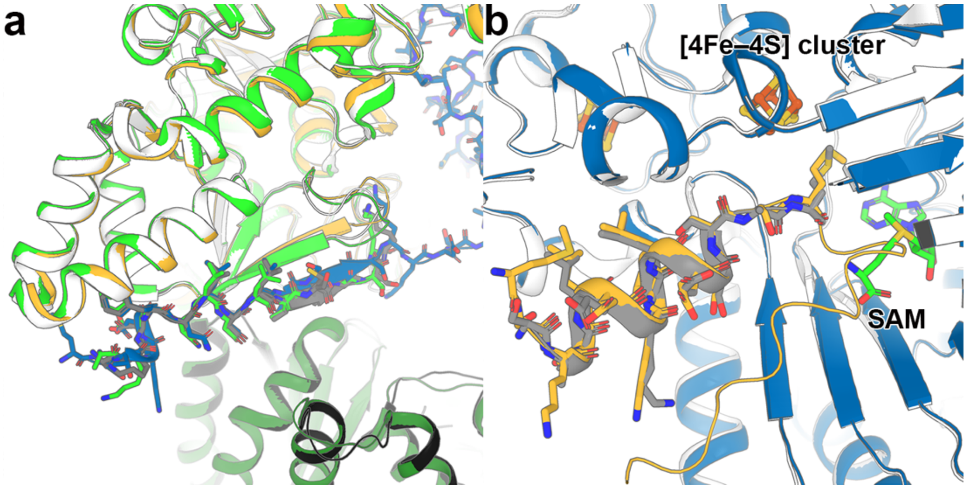


**Supplementary Fig. 2: Superimposition of predicted structures of RiPP precursor peptides and their tailoring enzyme complexes. a** Superimposition of the predicted NisA (blue)–NisB (yellow) model with the crystal structures (PDB IDs: 4WD9, green; and 6M7Y, white/gray/black). The predicted ipTM and ipSAE were 0.68 and 0.66, respectively. In 4WD9, the precursor peptide NisA is fused to NisB in the same chain. **b** Superimposition of the predicted StrA (orange)–StrB (blue) complex from *Streptococcus thermophilus* LMD-9 (BGC0001209) with the crystal structure of its homologous complex, SuiB (white)–SuiA (gray), from *Streptococcus suis* (PDB ID: 5V1T). The predicted ipTM and ipSAE were 0.82 and 0.79, respectively. SuiB and StrB share 94.98% sequence identity. The [4Fe–4S] cluster and (radical) SAM cofactor are shown as stick models.

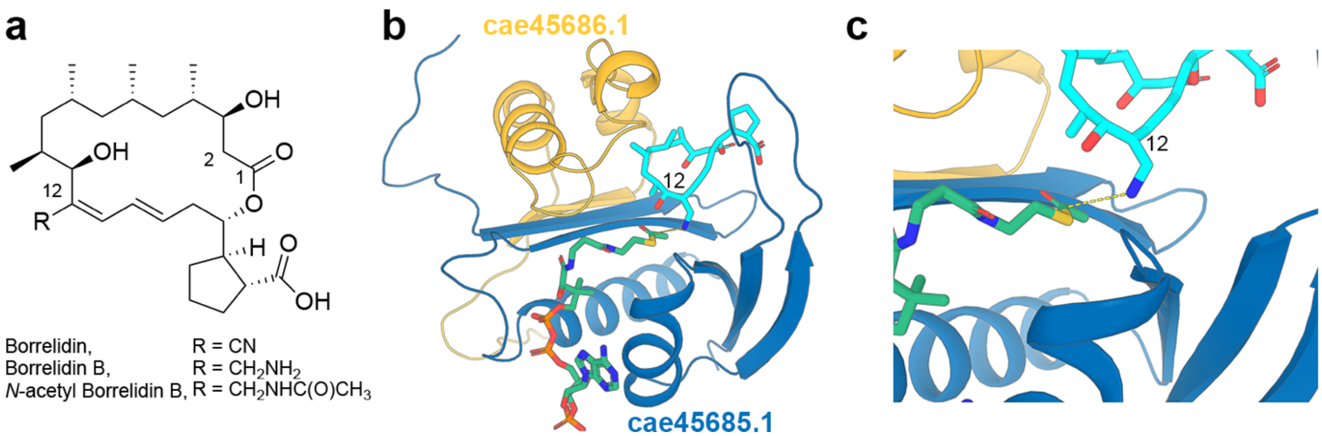


**Supplementary Fig. 3: Proposed catalytic mechanism of *N*-acetylborrelidin B. a** chemical structure of borrelidin, borrelidin B, and *N*-acetylborrelidin B. **b** Modeled cae45685.1 (blue)–cae45686.1 (yellow) heterodimer in complex with acetyl coenzyme A (green) and borrelidin B (cyan). **c** Close-up view of the two ligands. The dashed line is drawn between the putative reactive atoms.

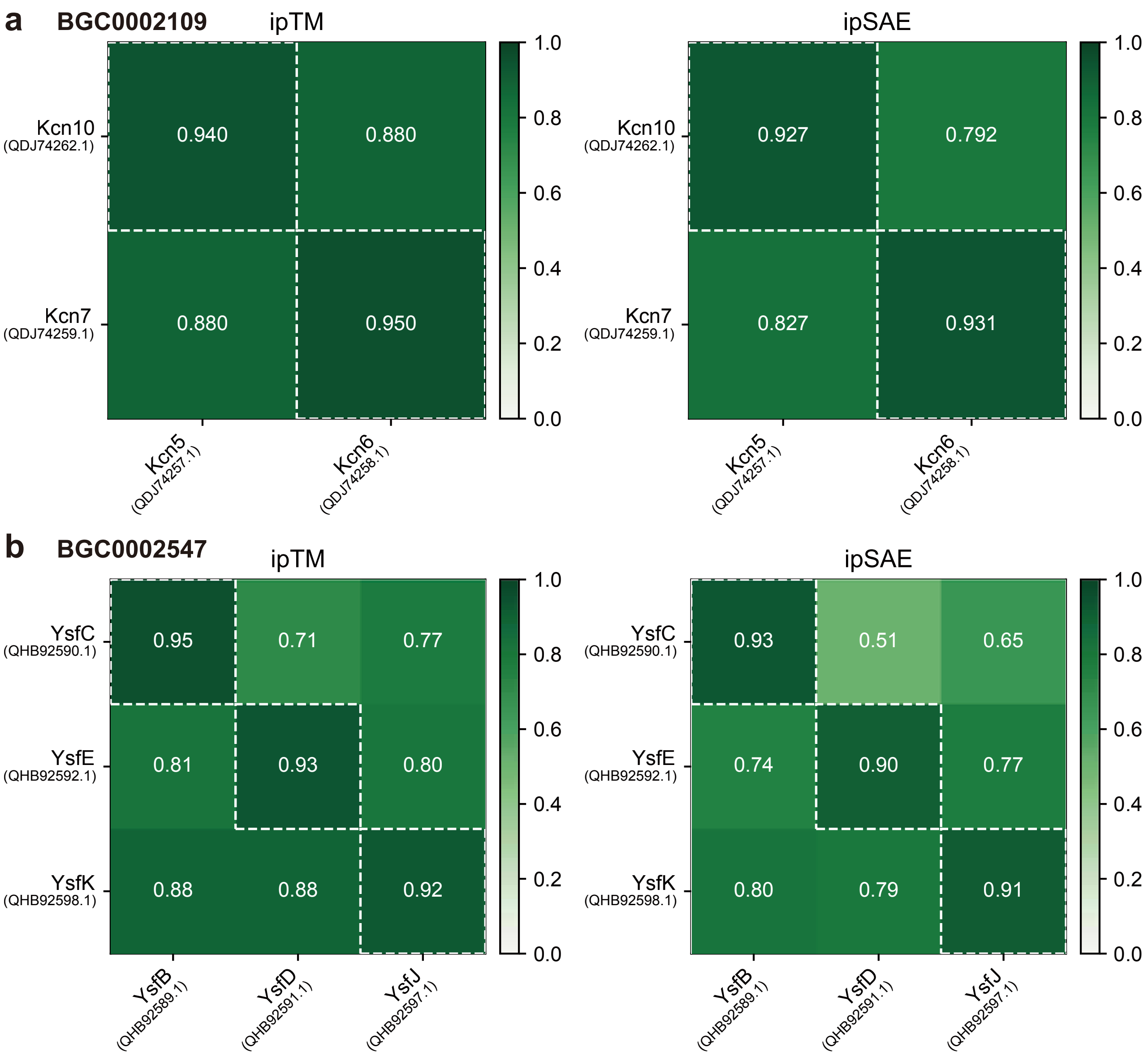


**Supplementary Fig. 4**: Computed ipTM and ipSAE values for complex prediction of the KS (horizontal) and CLF (vertical) proteins in the same BGC. **a** kitacinnamycin A BGC (BGC0002109)[6, 7]. **b** youssoufenes BGC (BGC0002547)[8]. The correct KS–CLF pairs identified by experiments are indicated by white dashed lines.

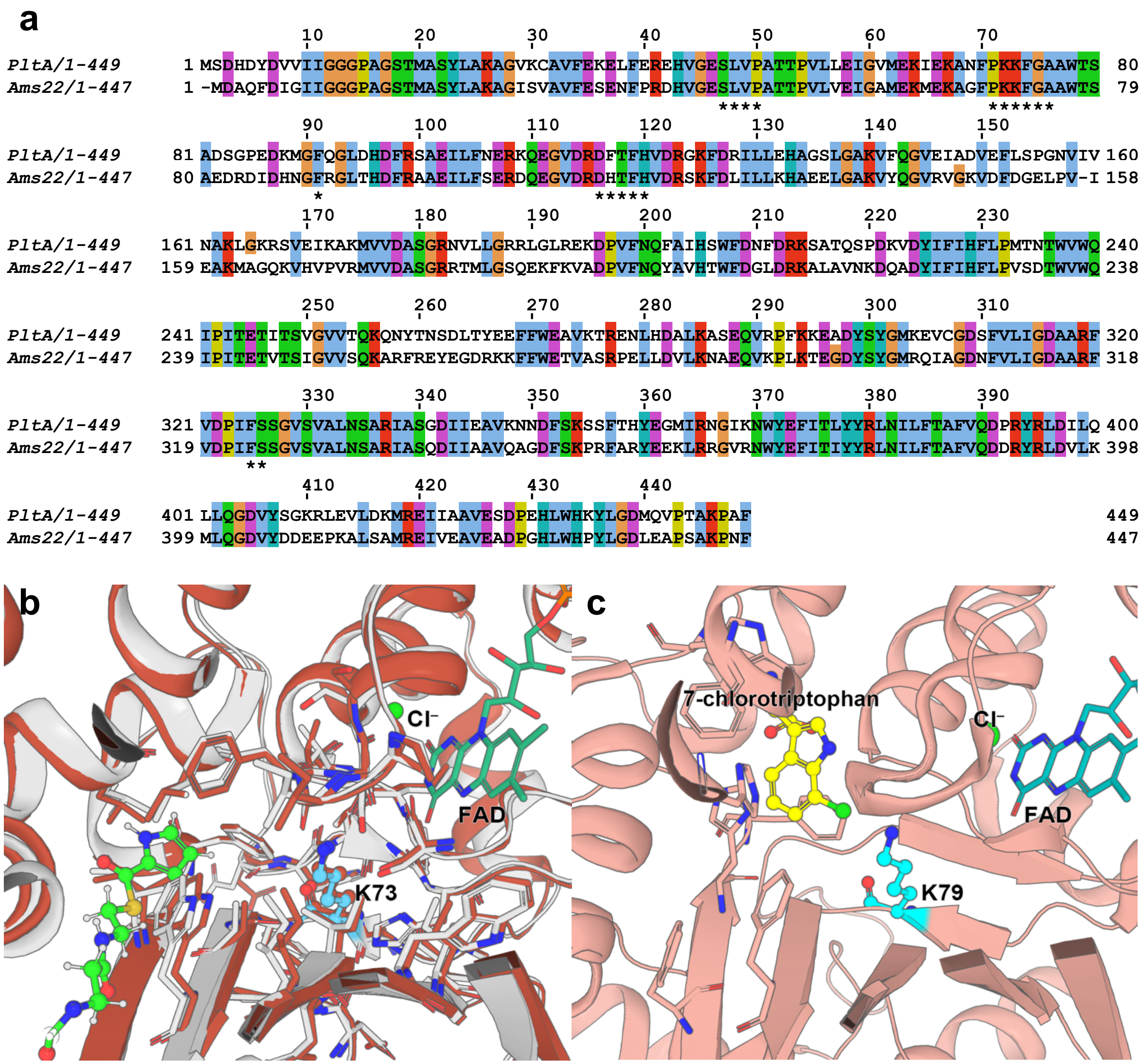


**Supplementary Fig. 5:** Comparison of PltA from the pyoluteorin BGC (BGC0000127) and Ams22 from the armeniaspirol BGC (GenBank ID: AZY92002.1). **a** Pairwise sequence alignment. Asterisks indicate residues located within 6 Å of the catalytic residue K73 of PltA. **b** Superimposition of the predicted structure of Ams22 (brown) onto the crystal structure of PltA (PDB ID: 5DBJ; white). Residues within 6 Å of the catalytic residue K73 of PltA are shown in stick representation. The modeled pyrrole substrate covalently attached to the phosphopantetheinyl arm is shown in yellow green. **c** Crystal structure of tryptophan 7-halogenase in complex with 7-chlorotryptophan (PDB ID: 2AR8)[9].

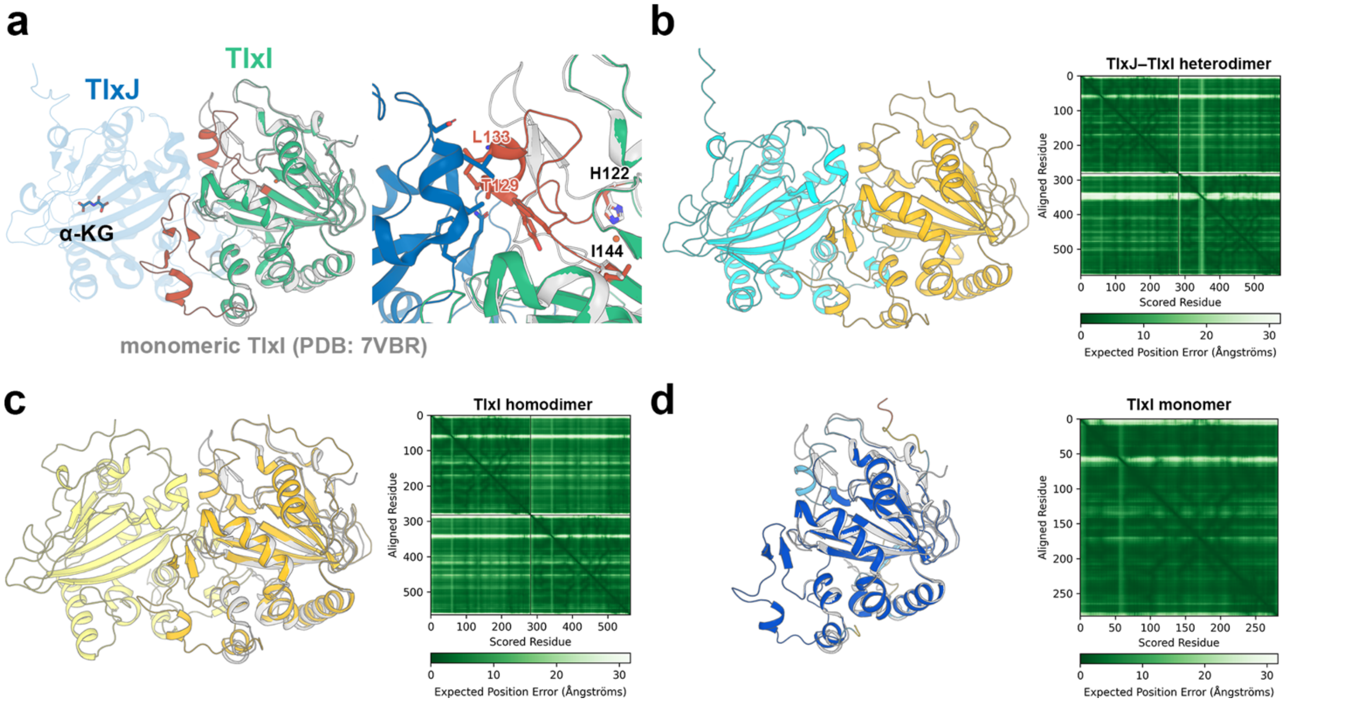


**Supplementary Fig. 6: Comparison of the crystal structure of TlxJ–TlxI and the predicted complex structure. a** Superimposition of the monomeric TlxI crystal structure (PDB ID: 7VBR) onto the heteromeric TlxJ–TlxI complex crystal structure (PDB ID: 7VBQ). Residues 122–144 of TlxI, which undergo large conformational changes, are highlighted in red. **b–d** Predicted models of the TlxJ–TlxI heterodimer (**b**), the TlxI homodimer (**c**), and the TlxI monomer (**d**). Their corresponding PAE matrices are shown in the right panels. In **c** and **d**, the crystal structure of TlxI is displayed in white for comparison. In **d**, TlxI is colored according to the pLDDT coloring (orange, 0–50; yellow, 50–70; cyan, 70–90; and blue, 90–100).
